# ProteinPrompt: a webserver for predicting protein-protein interactions

**DOI:** 10.1101/2021.09.03.458859

**Authors:** Sebastian Canzler, Markus Fischer, David Ulbricht, Nikola Ristic, Peter W. Hildebrand, René Staritzbichler

## Abstract

**Motivation:** Protein-protein interactions play an essential role in a great variety of cellular processes and are therefore of significant interest for the design of new therapeutic compounds as well as the identification of side-effects due to unexpected binding. Here, we present `ProteinPrompt`, a webserver that uses machine-learning algorithms to calculate specific, currently unknown protein-protein interactions. Our tool is designed to quickly and reliably predict contacts based on an input sequence in order to scan large sequence libraries for potential binding partners, with the goal to accelerate and assure the quality of the laborious process of drug target identification.

**Methods:** We collected and thoroughly filtered a comprehensive database of known contacts from several sources, which is available as download. `ProteinPrompt` provides two complementary search methods of similar accuracy for comparison and consensus building. The default method is a random forest algorithm that uses the auto-correlations of seven amino acid scales. Alternatively, a graph neural network implementation can be selected. Additionally, a consensus prediction is available. For each query sequence, potential binding partners are identified from a protein sequence database. The proteom of several organisms are available and can be searched for contacts.

**Results:** To evaluate the predictive power of the algorithms, we prepared a test dataset that was rigorously filtered for redundancy. No sequence pairs similar to the ones used for training were included in this dataset. With this challenging dataset, the random forest method achieved an accuracy rate of 0.88 and an area under curve of 0.95. The graph neural network achieved an accuracy rate of 0.86 using the same dataset. Since the underlying learning approaches are unrelated, comparing the results of random forest and graph neural networks reduces the likelihood of errors. The consensus reached an accuracy of 0.89. `ProteinPrompt` is available online at: http://proteinformatics.org/ProteinPrompt

The server makes it possible to scan the human proteome for potential binding partners of an input sequence within minutes. For local offline usage, we furthermore created a `ProteinPrompt` Docker image which allows for batch submission: https://gitlab.hzdr.de/Proteinprompt/ProteinPrompt. In conclusion, we offer a fast, accurate, easy-to-use online service for predicting binding partners from an input sequence.

## 1 Introduction

Protein interactions are key to the complex molecular interplays of cellular processes. The driving forces of these molecular networks are protein interactions rather than the individual functions of single protein components (Pawson, 2004). Biological processes, such as cellular organization, communication, immune responses, and the regulation of transcription and translation, function appropriately only when various proteins interact and work together properly.

Laboratory identification and validation of protein interactions often relies on expensive, time-consuming biochemical and biophysical assays, including ELISA, western blot, immunoprecipitation, Förster resonance energy transfer (FRET) or cross-linking approaches, florescence anisotropy, microscale thermophoresis (MST), surface plasmon resonance spectroscopy (SRP), high-throughput screening methods (e.g., phage display), or combinations thereof.

A reliable *in silico* method for predicting protein-protein interactions (PPI) would therefore shed more light on the details of biological pathways and pharmacological responses. Computations may complement and guide biochemical assays. However, explicit molecular dynamics or docking approaches require structural detail, which is often unavailable. Even if the structure of a protein is known, these methods are computationally expensive and therefore impractical for scanning huge libraries of candidates. Therefore, when structural insight is lacking or when speed is crucial, other methods are needed. Non-structure-based computational approaches for identifying potential PPIs generally use an extensive dataset of known protein-protein interactions, combined with information about cellular localization, amino acid sequences, or secondary structures.

These methods may include phylogenetic trees (Pazos and Valencia, 2001), phylogenetic profiles (Barker and Pagel, 2005; Hamp and Rost, 2015a), graph based approaches (Yang et al., 2020), support vector machines (Li et al., 2019), or networkbased approaches (Yook et al., 2004; Clauset et al., 2008), stacked autoencoders (Sun et al., 2017), as well as (recurrent) convolutional neural networks (Hashemifar et al., 2018; Chen et al., 2019). In recent years, distinct prediction methods have been combined, e.g. convolutional neural networks and feature-selecting rotation forests (Wang et al., 2019). Algorithms from language encoding (Yao et al., 2019) and principle component analysis (Kong et al., 2020) were used to derive feature vectors. Structural features were exploited: (Singh et al., 2010; Das and Chakrabarti, 2021). Overall, the field of biology has recently seen a massive increase in applications using deep learning (Ching et al., 2018). Nevertheless, different proteome-wide prediction methods have demonstrated that knowledge of the amino acid sequence alone may be sufficient to identify novel, functional protein-protein interactions (Martin et al., 2005; Shen et al., 2007). These methods usually rely on statistical learning algorithms. Due to its significant advantages, which include simplicity, rapidity, and generality, this method of prediction has become more and more common in recent years (Ofran and Rost, 2003; Betel et al., 2007; Liu et al., 2012; Perovic et al., 2017; Pan et al., 2010; Chen et al., 2020; Das and Chakrabarti, 2021). Precalculated databases are availabe online: `PrePPI` (Zhang et al., 2013), `ProfPPI` (Tran et al., 2018), `STRING` (Szklarczyk et al., 2011), or `PIPS` (McDowall et al., 2009). Several webservices offer a limited number of pairwise predictions: `PSOPIA` (Murakami and Mizuguchi, 2014), `iLoops` (Planas-Iglesias et al., 2013), `iFrag` (Garcia-Garcia et al., 2017). To our knowledge, no webserver currently offers scanning entire proteoms.

In this paper, we present an online sequencebased approach to predicting PPIs. The tool’s predictive power was boosted through rigorous finetuning of the key elements of machine learning, including dataset generation and feature vector design. Auto-correlation of hydrophobicities combined with a random forest (RF) machine-learning algorithm led to maximum accuracy. The quality and speed of this system make it a suitable high-throughput method for scanning sequence libraries. Furthermore, we achieved a comparable accuracy rate using a graph neural network (GNN). Since this approach has a completely different mathematical structure, we provide it as an option for the user on the server. Additionally, a consensus method is available.

Therefore, `ProteinPrompt` (**protein pr**ediction **o**f **m**atching **p**ar**t**ners) may serve as a reliable tool for identifying potential interaction partners from an entire proteom. It can thus be used to help identify the yet-unknown biological roles of many proteins and may contribute to identifying new therapeutic targets.

## 2 Materials and Methods

To maximize the system’s predictive power, it is pivotal to optimize all key elements, including the collection of training and testing data, the calculation of feature vectors, and the selection and finetuning of the machine-learning algorithm. Many varied approaches to all these steps were tested. Here, we focus on the approaches that resulted in the highest accuracy rates.

### 2.1 Collecting data points

In order to create comprehensive training and testing data, we tried to collect as many trustworthy PPI annotations as possible. We included data from various sources, such as the `Database of Human Interacting Proteins`^1^ (DIP) (Salwinski et al., 2004), the `Human Protein Reference Database`^2^ (HPRD) (Keshava Prasad et al., 2009), the `Protein Database`^3^ (PDB) (Berman et al., 2000), and the `Negatome Database`^4^ (Blohm et al., 2014). We also included annotations retrieved from the `KUPS`^5^ server (Chen et al., 2011), which mainly incorporates PPIs from `MINT`^6^ (Licata et al., 2012) and `IntAct`^7^ (Orchard et al., 2014). In addition to the negative annotations collected from the `Negatome Database`, the `KUPS` server generates negative data points based on the the following criteria: (1) the proteins are functionally dissimilar, (2) the proteins are located in different cellular compartments, and (3) the proteins are part of non-interacting domains.

After intense manual curation and after mapping the different names used to describe the same proteins, we derived a total list of 31,867 distinct human proteins. For this set, 73,681 positive PPIs were collected from the databases mentioned above. We then used `CD-hit` (Li and Godzik, 2006; Fu et al., 2012) to reduce this set to 41,482 positive protein-protein pairs with at most 50% sequence identity. For the negative pairs, we collected over 1.5 million unique protein-protein pairs, from which we randomly selected comparable number of PPIs to the size of the positive dataset, while still maintaining at most 50% sequence identity.

We separated the data into test and training data to estimate the quality of the optimized prediction model with an independent dataset, which was not involved in the training or similar to the data used for training. Our final training dataset contained 36,423 positive and 34,640 negative PPIs, while our testing dataset contained 5059 positive and 4817 negative data points.

A detailed description of the number of individual proteins in each data set can be found in the Supplementary Section 2.1.

To assess the prediction quality of `ProteinPrompt`, we compared our system to several other publicly available programs. As some of them have limited speed or upload capacity, it was not feasible to perform this test using our entire test dataset, which includes more than 10k data points. Instead, we randomly selected 470 positive and 470 negative pairs from our test set.

### 2.2 Analysis of data sets

In machine learning, data are often considered equal. Under this assumption, data can be divided into training and test data using random selection. This is clearly not the case for protein sequences, as the level of similarity among sequences may vary dramatically. (A comparison may be drawn from face recognition: Protein sequences can differ from one another as much as human faces differ from those of cats or whales.) Park et al. (Park and Marcotte, 2012) point out several combinatorial issues, which were further explored by Hamp et al (Hamp and Rost, 2015b). These issues are due to the pairwise nature of the input and should be considered when building the test dataset. It should be noted that our input data are sequence pairs that are identical in nature and therefore symmetric. This section addresses the relation between data points and their separation into test and training sets.

Obviously, it is significantly easier to predict binding partners for a query sequence that is very similar to one of the sequences in the training dataset than to predict binding partners for a sequence that has no similarity to any sequence in the training set. The latter case requires the algorithm to have “understood” some of principles that control the binding, while the former case only requires it to interpolate from known cases. The more predictions are based on actual understanding, the more general the results will be. However, it is very difficult to understand the complex, 3D interactions of macromolecules based on patterns in their sequences. A full understanding of these interactions requires some level of understanding of the folding of the individual proteins as well as their preferences regarding relative orientation.

Our goal was to train a method that was as general as possible, in the sense that it should not specialize in a certain class of human protein sequences. This meant that we needed to minimize redundancies in the datasets. Furthermore, we intended to perform a very rigorous test by reducing similarities between the training and testing data. As pointed out by Park et al., this approach may not lead to the best overall performance. However, it is the most rigorous way to test such a system.

Therefore, we analyzed the redundancies in our datasets by comparing similarities among data points. It should be noted that we were not looking for pairwise sequence similarity but the similarity of any given pair of sequences with any other pair of sequences. All datasets contain sequence pairs that are known to be binders or non-binders. First, we collected BLAST alignments for all the sequences in all datasets. We calculated similarity by dividing the number of identical positions by the length of the sequence. To compare the similarity of a pair A: (*A*_1_, *A*_2_) with pair B: (*B*_1_, *B*_2_) similarities were calculated in two possible combinations: Sim(*A*_1_, *B*_1_) with Sim(*A*_2_, *B*_2_) and Sim(*A*_1_, *B*_2_) with Sim(*A*_2_, *B*_1_). The combination resulting in a higher total similarity was selected. The 2D histogram in Figure 1 illustrates the low similarity between the binders in the training and test datasets. Thus, the selected test set represents a difficult challenge for the algorithm, and the accuracy rates reported here can be considered a worstcase estimate.

**Figure 1:**
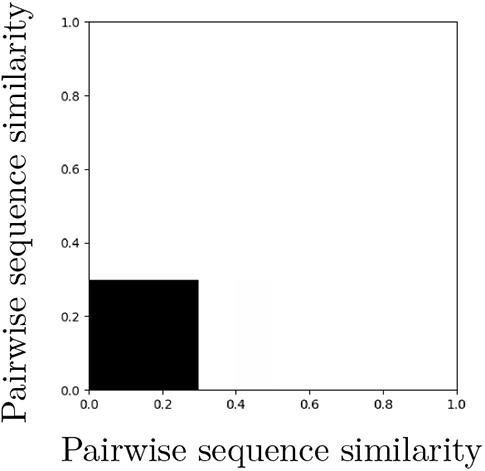
Redundancy analysis of binders in the training and test datasets. Depicted is a 2D histogram of similarity between sequence pairs known to be binders in the training and test datasets. The yellow area represents sequence pairs with no or very few similarities (below ~ 30%). It has a count of 1.8 · 10^8^. The violet area represents significantly smaller number of occurrences. A detailed description of this analysis is provided in section 2.2. The values of the histogram can be found in Table S1. Note that this figure is identical to the left image in Figure S1.

**Figure 2:**
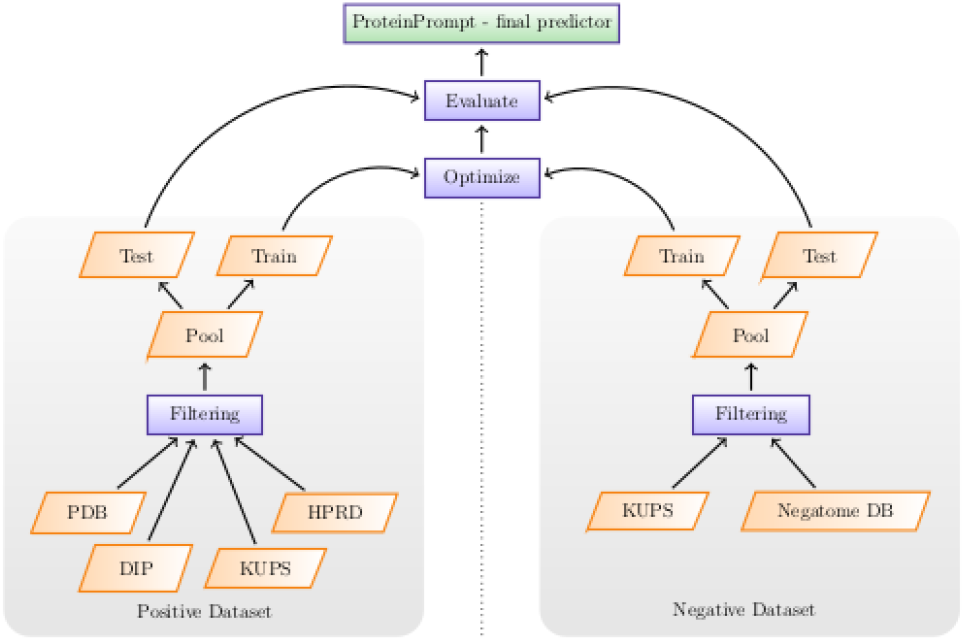
Overview of the training procedure.

Equivalent analyses with very similar results were performed on both datasets separately; the results are provided in the supplementary material (Figures S1, S2, S3). The values that are illustrated in the Figures are provided in Tables S1, S2, S3.

### 2.3 Selection of learning method

Many machine-learning algorithms are available. We used the `caret R` package (Kuhn, 2008) for fine-tuning and comparison. We tested several artificial neural network implementations, support vector machines, and tree-based methods. After determining that the RF approach performed best on our extensive testing and training datasets, we moved to a Python-based implementation to improve performance. A significant advantage of RFs is that overfitting is not a major problem.

We also extensively tested several more recent machine-learning methods, such as convolutional neural networks (CNN) and graph neural networks (GNN), which were implemented using the Tensorflow and PyTorch frameworks, respectively. CNNs were dismissed because of significant problems with overfitting. However, the GNN method resulted in comparable prediction quality to that achieved by the RF. Furthermore, the GNN method does not require any external preprocessing and can handle input data of varied sizes and structures. In our implementation, the GNN translates the data into fixed sized feature vectors, followed by a multilayer perceptron (MLP) that is predicting the binding. Feature vector calculation and predictions are trained in a single optimization scheme. Therefore, the GNN based implementation was trained on the raw data profiles.

### 2.4 Random forest classification

RF is a supervised learning algorithm that uses an ensemble of classification trees (Breiman, 2001). Each classification tree is built through bootstrap aggregation (bagging), a method of random sampling with replacement. From the original dataset, which contains *n* feature vectors, a random feature vector is selected *n* times and then copied to the dataset used to construct the tree. The copied feature vector is not removed from the original dataset. Thus, there will be multiple copies of some feature vectors. This means that different feature vectors are assigned varying degrees of importance. We defined a 420-dimension feature vector *F* = (*x*_1_, *x*_2_,…, *x*_420_) as the input for the RF model.

To construct an individual tree, a small random subset of a fixed size *m* ≪ 420 is extracted from the feature vectors at each node. The best split between these *m* values, which is the split that leads to the highest predictive power, is selected as the condition at each node. Each tree is grown as large as possible without pruning, resulting in low-bias trees. While this results in overfitting for a single decision tree, RF uses a high number of decision trees, which means that the algorithm has low variance. The final number of trees in the forest was set to 750 to optimize run time.

To further enhance the prediction quality of the RF, we also incorporated inverse and reverse PPI representations in our training. Each protein-protein pair (*A, B*) is therefore also described in the reversed direction (*B, A*) as well as two combinations with one inversed protein sequence: (*A,B_inv_*) and (*B,A_inv_*). For example *A*: ‘ANLMK’, and *A_inv_*: ‘KMLNA’. Among the eight possible protein pair representations, the combination of the four listed representations yielded the best prediction results and was therefore used in our RF model. This quadrupling of the data was not necessary for the GNN implementation.

#### 2.4.1 Feature vector calculation

Feature vector calculation includes extracting and transforming sequence-based information into a numerical vector of a constant size. Therefore, it is essential to extract the properties that direct the protein-protein interactions.

Each amino acid sequence of a protein-protein complex was transformed into a sequence of numerical values representing seven sequence-derived physicochemical properties. These properties are hydrophobicity (Eisenberg et al., 1984; Koehler et al., 2009), hydropyhilicity (Hopp and Woods, 1981), the volume of the side chains of amino acids (Krigbaum and Komoriya, 1979), polarity (Grantham, 1974), polarizability (Charton and Charton, 1982), solvent-accessible surface area (Rose et al., 1985), and the net charge index of the side chains of amino acids (Zhou et al., 2006) The properties were calculated for each residue in the sequence. These scales are commonly used for protein recognition (Ding and Dubchak, 2001) and to predict protein interactions (Bock and Gough, 2001, 2003), protein alignment (Stamm et al., 2013), protein structure (Durham et al., 2009), or protein functional families (Cai et al., 2003). These applications suggest that these properties significantly contribute to the stability of protein-protein complexes. Each amino acid scale was normalized as follows:

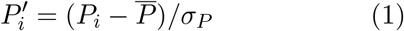

where 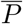 is the mean and *σ* is the standard deviation of the scale-based descriptor covering 20 amino acids, respectively:

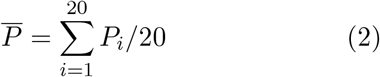

and

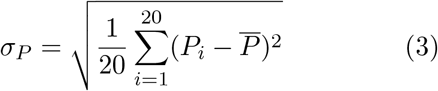

Auto-correlation (AC) was then used to transform the data into appropriate feature vectors as follows:

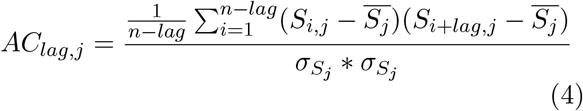

where *S_j_* is the translated amino acid sequence using the normalized scale-based descriptor 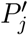 with *j* = 1, 2,…, 7, *n* is the length of sequence S, *lag* = 1, 2,…, 30 is the shift for which the autocorrelation is calculated, 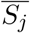 and *σ_S_j__* are the mean and standard deviation of the translated sequence, respectively. Ding *et al*. (2016) showed that a maximum *lag* of less than 30 tends to lose useful information, while larger values may induce noise. Accordingly, the number of AC values for each of the seven scales is 30. The feature vector describing any individual amino acid sequence has 7 · 30 = 210 elements or dimensions. Thus, for a pair of sequences, the feature space has 420 dimensions.

### 2.5 Graph neural networks

Graph neural networks (GNN) are a relatively new type of neural networks that operate on graphs (Scarselli et al., 2009), (Battaglia et al., 2018). Features can be assigned and predicted on a node, edge, or graph level. The algorithm takes a graph as input, performs computations on the graph itself through a process called message passing, and returns a graph of identical structure but updated features.

In a message passing round, first, the edge features are updated by a learnable update function that takes connected nodes and current edge features into account. Afterwards, another learnable function calculates new node features based on the aggregation of all connected edges and current node features. In the final step, a graph-level target is constructed by a third learnable function that takes an aggregation of all the nodes and edges as input. Through this process, every node and edge collects information about its local region in the graph; these data are used to infer global features.

We used GNNs to condense the information in protein sequences of varying lengths to a fixed-sized vector representation of each sequence. One graph is constructed for each sequence; nodes represent amino acids, and edges are constructed between the nodes of adjacent amino acids. The node features were encoded using a MLP that transformed the values of the amino acid profiles into a 32-dimension vector. The edge features were encoded from a vector of same size that was initialized with ones. The update functions for the nodes and edges and for the graph were implemented by MLPs.

After five rounds of message passing, a final aggregation function is performed on the graph to calculate a 128-dimension graph-level feature from all nodes and edges. This output can be understood as an abstract representation of each protein. This is done for both sequences. Then, their feature vectors are concatenated to one 256-dimension vector, which is then used as the input for the MLP. The last network returns the prediction of the binding propensity of the two proteins.

The model was built using PyTorch and the PyTorch geometric library. The message passing graph net block was realized using its MetaLayer class with an additional edge update on the graph. Parameters were initialized as described by (Glorot and Bengio, 2010), and a leaky rectified linear unit (ReLU) (Maas et al., 2013) was used for activations, except in the final layer of the last MLP, where a sigmoid activation was used. Training was done using binary cross-entropy loss and the AdamW (Loshchilov and Hutter, 2017) optimizer with a learning rate of 1e-5 for 350 epochs. The iteration with the best performance on the test set was refined over an additional 50 epochs with a learning rate of 1e-7. See Figure S4 in the supplementary materials for a graphical illustration of our implementation.

### 2.6 Consensus of RF and GNN

We trained a simple neural network with the outputs of RF and GNN for performing a consensus prediction. The scores returned by RF and GNN show rather different distributions. Thus, a linear combination is unlikely to result in the best possible consensus. We selected a NN with two inputs, two hidden layers with 8 neurons and a single output neuron (2-8-8-1).

## 3 Results

### 3.1 Performance RF

Based on our test dataset of 5094 positive and 5094 negative cases (binders vs. non-binders), we plotted the receiver operation characteristics (ROC) curve, shown in Figure 3. This allowed us to estimate the overall quality of the predictions and also provides a visual overview of the relationship between true and false positives. Each point on the curve shows how many falsely predicted binders should be expected for a given number of correctly predicted ones. As the datapoints are sorted by their predicted binding propensity, a reasonable threshold can be selected. This allows to fine-tune the balance between sensitivity and specificity.

**Figure 3:**
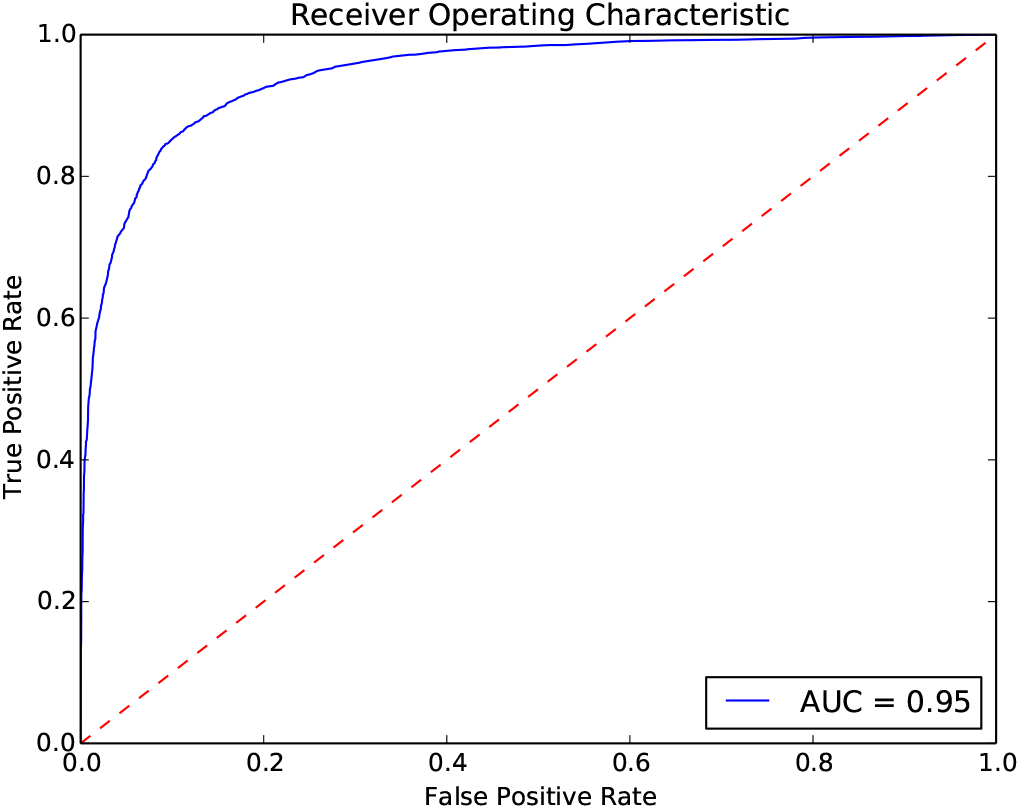
Performance of `ProteinPrompt` on the entire test dataset of over 10,000 PPIs.

Sensitivity is the ratio of correctly predicted binders (true positives) to actual binders. Specificity is the ratio of correctly predicted non-binders (true negatives) to all non-binders. For example, in Figure 3, a false positive rate of 0.03 will result in a true positive rate of ~0.6. Thus, selecting this point (its associated score) as the threshold will result in a hit rate of 60% and only 3% false predictions should be expected.

An ideal signal would lead to a rectangular plot with an area under the curve (AUC) of 1. The other extreme, pure noise, would result in a diagonal line with an AUC of 0.5. Our method results in an AUC of 0.95, specificity of 0.88, sensitivity of 0.87, and an accuracy rate of 0.88. Currently, our method balances specificity and sensitivity to avoid unwanted bias in different applications.

#### 3.1.1 RF compared to other tools

We compared our optimized RF system to publicly available tools such as `SPPS` (Liu et al., 2012), `TRI_tool` (Perovic et al., 2017), and `LR_PPI` (Pan et al., 2010). Due to limited access, we used the reduced test dataset, as described in the methods section. We also re-evaluated `ProteinPrompt` on the reduced test dataset to ensure comparable results. The results of our evaluation using the reduced test dataset are somewhat different from those we obtained using the entire test dataset.

The results shown in Table 1 and the ROC plot in Figure 4 suggest that `ProteinPrompt` outperforms the other three methods. Furthermore, `ProteinPrompt` balances sensitivity and specificity. `SPPS` and `LR_PPI` have excellent sensitivity of 0.96 and 0.91; however, their specificity is rather poor: 0.34 and 0.16, respectively. `TRI_tool` on the other hand, shows a massive bias towards specificity, which is 0.95, compared to a sensitivity of 0.32. Furthermore, the overall prediction accuracy and the AUC of our tool are significantly higher. Accuracy: 0.86 versus 0.66 (`SPPS`), 0.63 (`TRI_tool`) or 0.53 (`LR_PPI`). AUC: 0.94 versus 0.77 (`SPPS`), 0.73 (`TRI_tool`), and 0.60 (`LR_PPI`), respectively.

**Figure 4:**
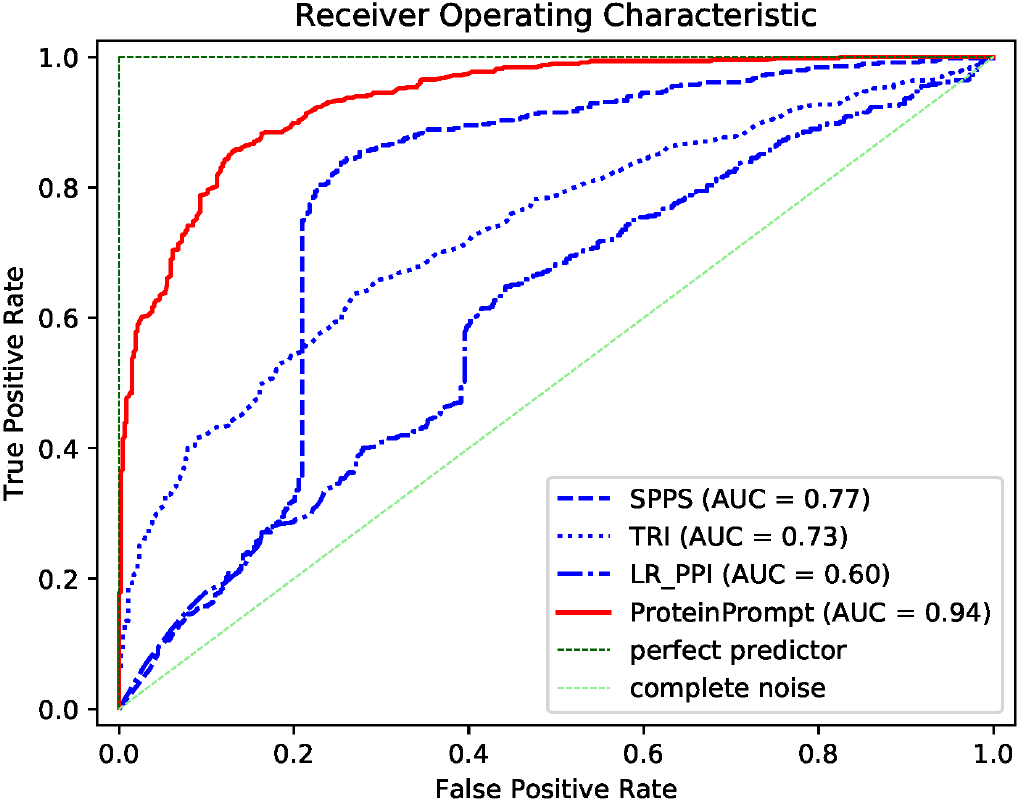
ROC curves for `ProteinPrompt, SPPS, TRI_tool, LR_PPI` using the reduced test dataset of 968 protein-protein pairs.

**Table 1:** Quality measures of `ProteinPrompt`compared to other publicly available tools, such as `SPPS` (Liu et al., 2012), `TRI_tool` (Perovic et al., 2017), and `LR_PPI` (Pan et al., 2010). AUC, specificity, sensitivity, and accuracy are listed.

| Tool | AUC | Spec. | Sens. | Acc. |
| --- | --- | --- | --- | --- |
| **ProteinPrompt** | 0.94 | 0.88 | 0.84 | 0.86 |
| **SPPS** | 0.77 | 0.34 | 0.96 | 0.66 |
| **TRI.tool** | 0.73 | 0.95 | 0.32 | 0.63 |
| **LR.PPI** | 0.60 | 0.16 | 0.91 | 0.53 |

#### 3.1.2 Individual test cases

We further tested the detection rate of `ProteinPrompt` using experimentally verified protein interaction partners (EV PPIs). We compared the output of `ProteinPrompt` to that of the STRING database^8^, which has been shown to include a very high number of experimentally proven PPIs (Bajpai et al., 2020). Here, the EV PPIs of five different prominent proteins with various cellular functions (Table S4 in the supplementary material) with high confidence (score ¿ 0.7) were investigated. The PPI predictions of `ProteinPrompt` were compared to those of the STRING database. `ProteinPrompt` found all but one of the EV PPIs output by the STRING Database; for the SRC gene one of nine binding partners was not identified (Table S4). The scores given by the STRING database and the scores of `ProteinPrompt` were then statistically analyzed and plotted as boxplots to enable direct comparison (Figure 5). On average, `ProteinPrompt` predicted all identified EV PPIs with comparable or better accuracy than the STRING database (Figure 5, Table S4). However, due to the complexity of PPIs in nature, STRING scores do not necessarily reflect real protein-protein affinities. On our test dataset, `ProteinPrompt` was able to rapidly predict EV PPIs with an average accuracy rate of 0.89 and a SD of 0.09. From a userâĂŹs perspective, `ProteinPrompt` identified 98% of all EV PPIs with a predicted score above 0.7, 84% of all EV PPIs with a predicted score above 0.8, and 46% of all EV PPIs with a predicted score above 0.9.

**Figure 5:**
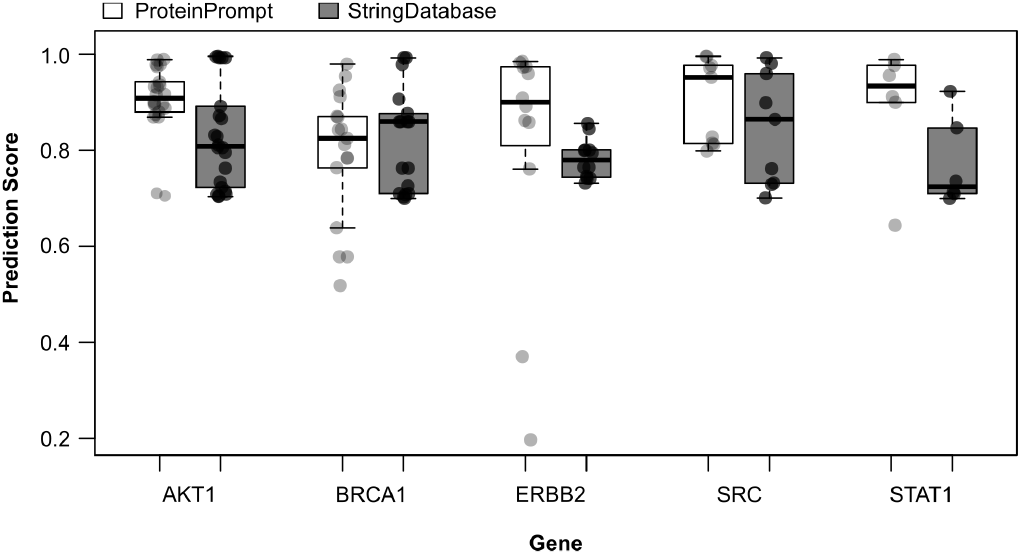
EV PPI scores from `ProteinPrompt` (green) and from STRING database (grey) for a test set of five different proteins, indicated by their gene names in the boxplot.

#### 3.1.3 Dataset of Park and Marcotte

As an additional test, we used the Park and Marcotte (2012) dataset on which a number of methods were tested, allowing a direct comparison. Park and Marcotte (2012) tested 7 methods, Ding *et al*. (2016) extended this with an additional 3 methods. `ProteinPrompt` in its current form performs similarly on this dataset as it does on our own test data. For all human subsets, `ProteinPrompt` achieved significantly higher accuracy than all other methods. When `ProteinPrompt` was trained with this data, the performance is similar to the other top ranking tools. For most human subsets, the retrained version of `ProteinPrompt` ranked second. First, this confirms that combining RF with autocorrelation has the potential to achieve high accuracy. Second, this test highlights the critical impact of the dataset on the final performance. For the yeast data, `ProteinPrompt` does not perform as well, indicating limited transferability to other organisms, a fact also observed with other learning methods. Most learning methods are only valid within the domain of their training data. A detailed description can be found in the Supplementary Materials.

### 3.2 Performance GNN

In the first test, we used the same seven amino acid scales that were used for the RF. We compared this to a graph topology in which 13 and 15 scales were assigned to each amino acid. The resulting accuracy rates are listed in Table 2.

**Table 2:** Accuracies of GNNs trained on different numbers of amino acid profiles.

| # profiles | Accuracy |
| --- | --- |
| 7 | 83.5% |
| 13 | 86.0% |
| 15 | 83.8% |

Adding as many residue scales as possible to the graph does not improve the accuracy. Currently, the maximum accuracy is 86% for 13 residue scales. Extensive future research would be needed to further maximize the performance of the GNN.

In cross validations, in which 20% of the training data were used as validation set, the accuracies for the validation data were nearly identical to the one of the test data. This reflects the low level of redundancy within the training dataset, which is shown in Figure S3. We therefore considered it acceptable to perform a final optimization with 95% of the test dataset added to the training data. The resulting model is the one uploaded to the server. This strategy should lead to the most generally applicable model.

### 3.3 Performance consensus RF and GNN

The consensus using the RF and GNN as input was leading to a slight improvement with a final accuracy of 89.4%.

### 3.4 Webserver implementation

`ProteinPrompt` is available online as a webserver at: http://proteinformatics.org/ProteinPrompt Basic usage is as simple as providing an input sequence. By default, `ProteinPrompt` will search our manually curated database of human proteins, which contains 27,223 sequences. Scanning the entire human database takes approximately one minute for the default method RF. For GNN the search is even faster. The consensus has to execute both methods and therefore requires correspondingly more time. Other, more extensive databases are also provided, including mammalian, vertebrate, and metazoan protein sets. Searching these databases takes considerably longer. For example, the vertebrate database has 91,592 entries; therefore, searching it takes approximately three times longer than searching the human protein database.

The server is free for academic users. Providing an email address is optional. `ProteinPrompt` was originally optimized for sequences with a minimum length of 16 AA. The sequence length of the uploaded proteins is detected automatically. When the user provides a sequence shorter than 16 AA, a warning appears, but the calculation is still performed. As output, a ranked and scored list of proteins from the database is returned; this list can be downloaded.

## 4 Discussion

`ProteinPrompt` offers a reliable, fast way to predict protein interaction partners based on protein sequences. It is available as an easy-to-use online tool and is thus accessible to non-expert users. In order to develop this fast, reliable service, we optimized the learning algorithm, the binding database, and the representations of the sequences. We determined that the RF algorithm combined with autocorrelation on seven amino acid scales resulted in the highest accuracy.

We also determined that the GNN method performed nearly as well as the RF algorithm. To support consensus building, we offer both implementations on our server. It is remarkable that the RF algorithm, which is conceptually comparably simple, performs so well on such a complex task. This is even more remarkable considering that the RF approach, unlike the GNN, does not include simultaneous optimization of feature vector creation and model building.

An extensive database with limited redundancy was essential to reliably test the tool. This database was obtained through several iterations of manual curation of the test and training datasets. Despite strict separation of the training and test datasets, which posed a significant learning challenge, `ProteinPrompt` turned out to perform very well compared to other available servers and methods. `ProteinPrompt` is reasonably accurate and can scan the entire human proteome within approximately one minute. To our knowledge, `ProteinPrompt` is currently the fastest online service available for scanning different proteoms to identify potential binding partners based on a sequence level. It is reasonable to assume that expanding the training data would lead to higher accuracy rates. Based on our extensive tests, we expect `ProteinPrompt` to support a better understanding of the complex networks of proteinprotein interactions, which are the basis for a broad range of biological mechanisms.

## Acknowledgements

We extend our thanks to Johanna Tiemann, Alexander Vogel, Thorsten Kaiser and Daniel Wiegreffe for proofreading.

## Funding

This work was funded by the Sächsische Aufbaubank (SAB) [100256197, 100267985], and Deutsche Forschungsgemeinschaft (DFG, German Research Foundation) through CRC 1423 [421152132; sub-projects Z04]

## Footnotes

1 http://dip.doe-mbi.ucla.edu/

2 http://www.hprd.org/

3 www.rcsb.org

4 http://mips.helmholtz-muenchen.de/proj/ppi/negatome

5 http://www.ittc.ku.edu/chenlab/kups/

6 https://mint.bio.uniroma2.it/

7 https://www.ebi.ac.uk/intact/

8 https://string-db.org

## Notes

### Competing Interest Statement

The authors have declared no competing interest.

