## Supplementary Material for "ProteinPrompt: a webserver for predicting protein-protein interactions"

### ProteinPrompt: a webserver for the prediction of protein-protein-interactions - Supplementary material

December 15, 2021

Note: the numbering of the sections in this document is identical to the numbering of the sections in the main manuscript to which the sections here refer.

#### 2.1 Collecting data points

In total, we collected 41,482 positive protein-protein pairs with at most 50% sequence identity, while collecting 39,457 negative protein-protein pairs. The positive set contains 10,538 individual proteins and the negative set contains 13,304 individual proteins. A summary of the respective test and training sets can be seen in Table S1. An overview of the overlapping proteins among those four datasets is depicted in Figure S1. The following sections investigate the datasets in further detail.

| Dataset | # protein pairs | # proteins |
|---|---|---|
| pos. training | 36423 | 10175 |
| neg. training | 34640 | 13264 |
| pos. testing | 5059 | 4591 |
| neg. testing | 4817 | 6816 |

Table S1: Summary of individual protein pairs and number of unique proteins per dataset.

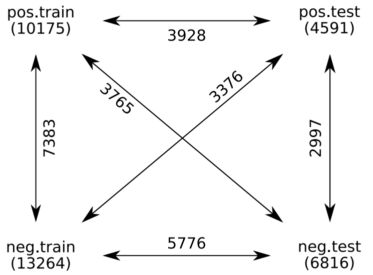


Figure S1: Number of overlapping proteins between different datasets. Numbers in parenthesis denote the number of unique proteins in each dataset.

#### 2.2 Analysis of the data sets

In this section, we examine the relationship of sequence pairs in our datasets rather than of individual sequences as in the previous section. Because of the pairwise nature of binding, it is critical that different datasets do not contain similar pairs. A BLAST search was performed with an e-value cutoff of 100, creating pairwise alignments for all sequences in the datasets. Alignments with e-values larger than the threshold are considered insignificant or unrelated. Sequence pairs for which alignments were returned had similarities above 30%. Thus, in our depiction unaligned sequences with an e-value above 100 are subsumed into the region with similarity up to 30%. As we cannot distinguish anything below 30% similarity, we fused this into a larger bin. Above 30% there is a binning of 10%. However, in the range of 30 to 40% only few sequence pairs were aligned. Thus, we would consider this bin as rather unreliable.

Figure S2 shows the similarities between the training and test data, Figure S3 shows similarities between entries within the training dataset. Figure S4 shows similarities between entries of the test dataset. Tables S2, S3, and S4 show the data that has been visualized in the those three figures, respectively.

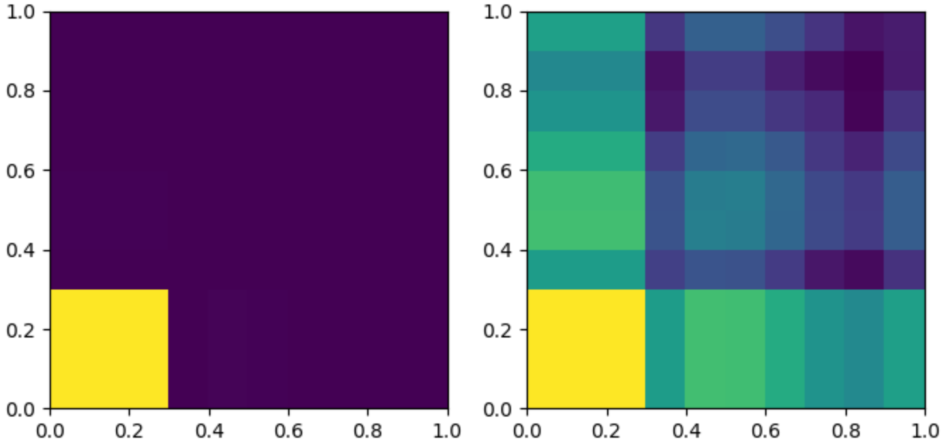


Figure S2: Redundancy analysis between binders in training and test data. Depicted is a 2D histogram of similarity between sequence pairs from binders of the training versus binding sequence pairs from the test dataset. The image on the right is in log-scale to visualize the background. See Table S2 for values. The image on the left is identical to Figure 1 in the main manuscript, except for the color scheme.

| | 0 - 30% | 30-40% | 40-50% | 50-60% | 60-70% | 70-80% | 80-90% | 90-100% |
|---|---|---|---|---|---|---|---|---|
| 0 - 30% | 177334846 | 126587 | 1405003 | 1315862 | 347567 | 73347 | 35897 | 152264 |
| 30-40% | 124375 | 394 | 1192 | 1117 | 312 | 51 | 29 | 192 |
| 40-50% | 1363827 | 1160 | 18244 | 15095 | 3560 | 715 | 319 | 1999 |
| 50-60% | 1245961 | 1059 | 14776 | 17522 | 4001 | 672 | 279 | 2034 |
| 60-70% | 337746 | 335 | 3555 | 4348 | 1566 | 251 | 88 | 716 |
| 70-80% | 75261 | 54 | 757 | 727 | 242 | 118 | 22 | 195 |
| 80-90% | 33803 | 36 | 332 | 330 | 72 | 30 | 19 | 61 |
| 90-100% | 161494 | 234 | 2453 | 2410 | 827 | 212 | 45 | 65 |

Table S2: Training versus test data for binders.

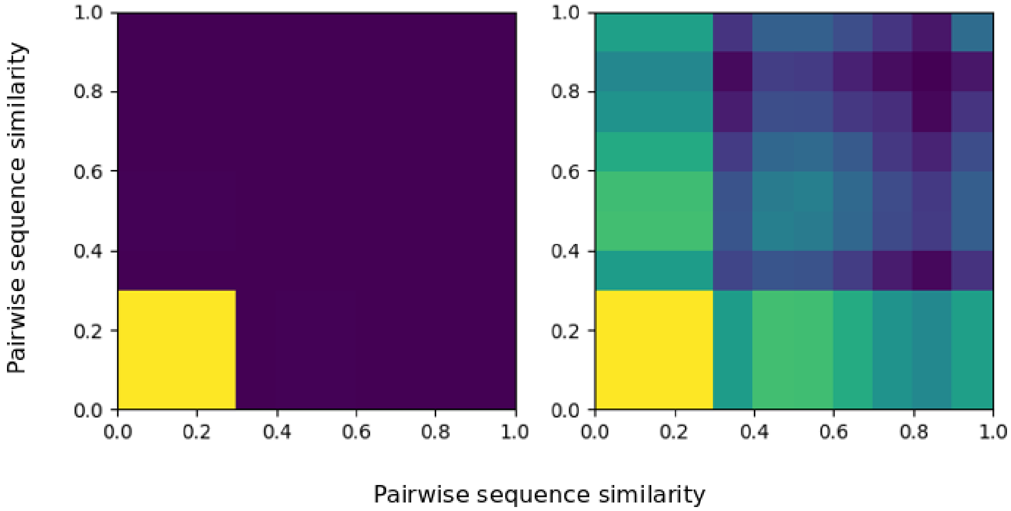


Figure S3: Redundancy analysis between binders in training data. See Table S3 for values.

| | 0 - 30% | 30-40% | 40-50% | 50-60% | 60-70% | 70-80% | 80-90% | 90-100% |
|---|---|---|---|---|---|---|---|---|
| 0 - 30% | 1277793513 | 924217 | 9843509 | 9199670 | 2460137 | 521258 | 241930 | 1074056 |
| 30-40% | 902712 | 3517 | 9117 | 8221 | 2330 | 464 | 193 | 1437 |
| 40-50% | 9548046 | 9517 | 126468 | 102733 | 26506 | 5021 | 2123 | 15149 |
| 50-60% | 8703841 | 8038 | 101608 | 126427 | 30691 | 5293 | 1955 | 15293 |
| 60-70% | 2375120 | 2188 | 27561 | 32355 | 12064 | 1846 | 611 | 5691 |
| 70-80% | 523609 | 476 | 5799 | 5629 | 1821 | 1107 | 178 | 1495 |
| 80-90% | 227925 | 201 | 2394 | 2153 | 565 | 224 | 133 | 357 |
| 90-100% | 1143738 | 1539 | 17066 | 17158 | 5817 | 1622 | 353 | 36939 |

Table S3: Training data cross similarities for binders.

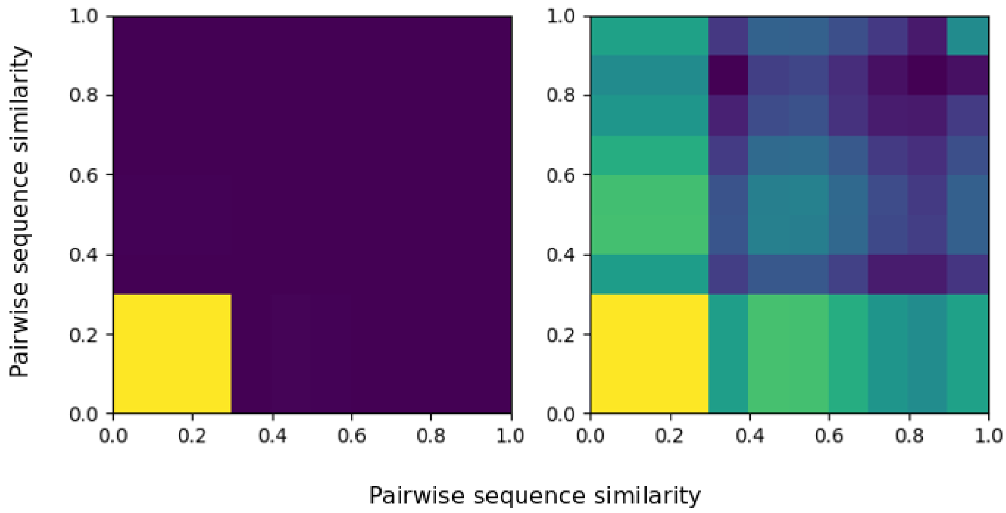


Figure S4: Redundancy analysis between binders in test data. See Table S4 for values.

| | 0 - 30% | 30-40% | 40-50% | 50-60% | 60-70% | 70-80% | 80-90% | 90-100% |
|---|---|---|---|---|---|---|---|---|
| 0 - 30% | 24618884 | 17412 | 199895 | 188349 | 48543 | 9988 | 5104 | 21649 |
| 30-40% | 16773 | 39 | 172 | 164 | 46 | 7 | 7 | 25 |
| 40-50% | 185493 | 145 | 2320 | 1977 | 463 | 73 | 40 | 280 |
| 50-60% | 175907 | 129 | 2127 | 2388 | 485 | 84 | 33 | 280 |
| 60-70% | 48105 | 35 | 516 | 594 | 182 | 32 | 18 | 104 |
| 70-80% | 10724 | 9 | 84 | 120 | 21 | 7 | 6 | 35 |
| 80-90% | 5011 | 2 | 41 | 59 | 16 | 4 | 2 | 4 |
| 90-100% | 22613 | 29 | 311 | 307 | 101 | 31 | 6 | 5071 |

Table S4: Testing data cross similarities for binders.

| Gene name | AKT1 | BRCA1 | ERBB2 | SRC | STAT1 |
|---|---|---|---|---|---|
| Protein | RAC-alpha serine/threonine-protein kinase | Breast cancer type 1 susceptibility protein | Receptor tyrosine-protein kinase erbB-2 | Proto-oncogene tyrosine-protein kinase Src | Signal transducer and activator of transcription 1-alpha/beta |
| Uniprot ID | P31749 | P38398 | P04626 | P12931 | P42224 |
| Prominent functions | Signal transduction, cell survival | DNA repair, tumor marker | Growth factor receptor, cell proliferation | Proto-oncogene, cell survival | transcription factor, immune response |
| Cell compartment | cytosol | nucleus | plasma membrane | cytosol | cytosol, nucleus |
| Number of EV PPIs in STRING.db (high confidence) | 22 | 17 | 12 | 9 | 6 |
| STRING.db score (Mean $\pm$ SD) | 0.829 $\pm$ 0.108 | 0.822 $\pm$ 0.107 | 0.783 $\pm$ 0.041 | 0.847 $\pm$ 0.118 | 0.772 $\pm$ 0.092 |
| Number of EV PPIs found with ProteinPrompt | 22 | 17 | 12 | 8 | 6 |
| ProteinPrompt score (Mean $\pm$ SD) | 0.903 $\pm$ 0.074 | 0.793 $\pm$ 0.138 | 0.811 $\pm$ 0.257 | 0.905 $\pm$ 0.088 | 0.896 $\pm$ 0.129 |

Table S5: Detailed comparison of the STRING database with ProteinPrompt.

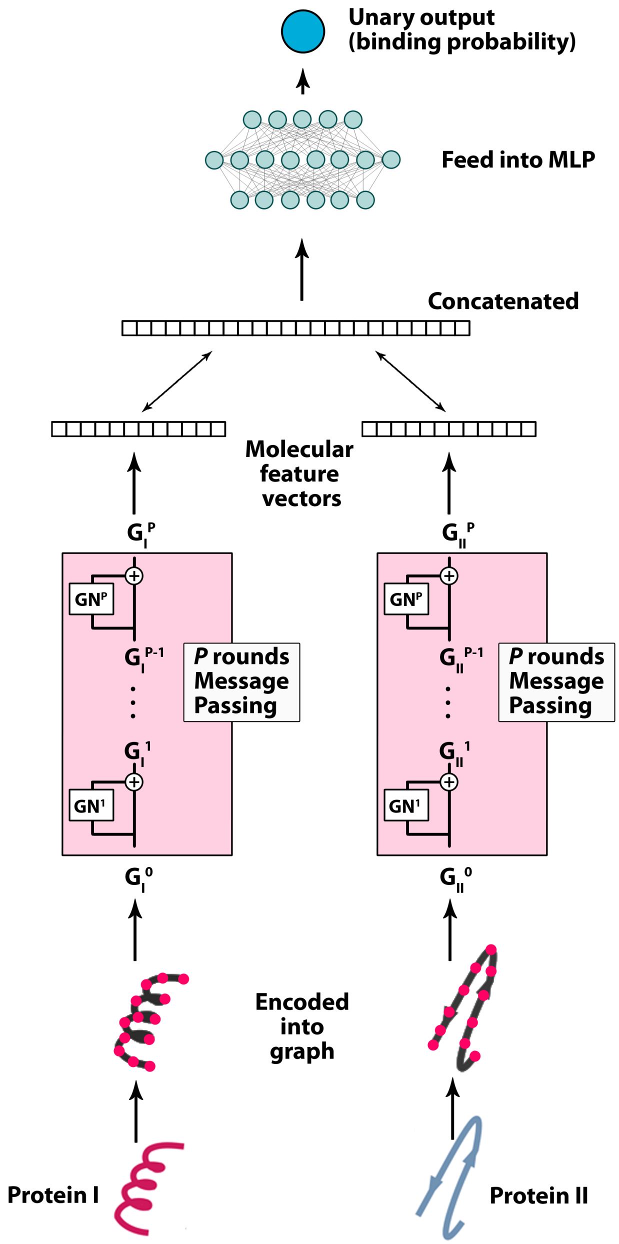


Figure S5: Schematic description of the graph neural network implementation. The proteins are encoded into graphs in which each amino acid is represented by one node and subsequent residues are connected by an edge. During the optimization procedure, the GraphNet (GN) modifies the aggregate and update functions. From each GNN, a feature vector is created. These feature vectors are concatenated and then passed to the final MLP, which predicts the binding. All elements of the entire scheme are optimized commonly in a single back-propagation.

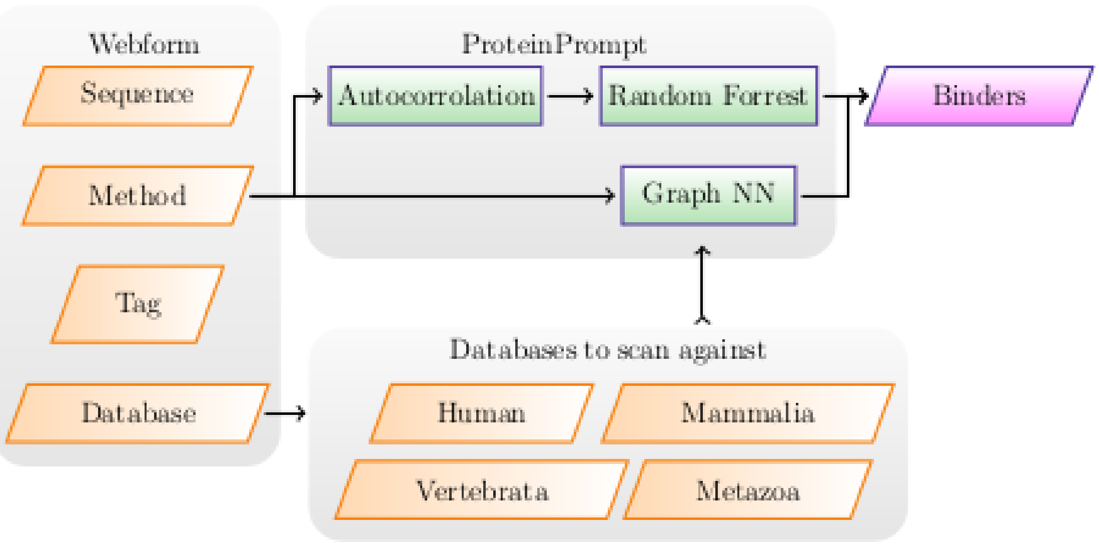


Figure S6: Server workflow. The user pastes the query sequence, provides a unique tag, and selects a method and organism to scan. The server returns a sorted table of potential binders.

##### 3.1.3 Verification of ProteinPrompt::RF with the datasets of Park and Marcotte

Park & Marcotte [1] performed a detailed analysis of the pairwise input of sequence based protein-protein interaction (PPI) predictions. As a result, the Marcotte lab provides the download of several training and test datasets [3]. In principle, they provide two human and two yeast datasets that differ in their composition: 'human_balanced', 'human_random', 'yeast_balanced', and 'yeast_random'. Each dataset contains 'typical' cross-validation sets (CV) but also sets where the test data was split into distinct classes (C1-C3), depending on the appearance of proteins in the respective training data, see Table S6 for a definition of those test classes. Park & Marcotte also provide the results of 7 different PPI predictions methods, which were trained on that data (M1-M7). Ding *et al* extended this by another 3 methods (MMI, NMBAC, MMI+NMBAC) [2]. The availability of this data has two major benefits: First, when developing a new method for PPI prediction, one can compare directly with the results of these 10 methods without having to go through the often time-consuming process of installing the software and all the necessary dependencies. Second, as in our case, having obtained results with higher accuracy than previous methods it is possible to distinguish whether this increase in prediction quality is due to an improved learning method or whether a more suitable dataset is the cause of the increase.

We performed two tests on each of the four available datasets. First, we ran the normal version of ProteinPrompt, trained on our human training data as described in the main text. The results for the test data in both human subsets 'random' and 'balanced' show a similar level of accuracy than on our own test data. However, it has to be noted that this is not a very robust result, as there may be significant overlap between our training data and the test data provided by Park & Marcotte. Since we have filtered our data very rigorously, as described in the previous section, the verification using our own test data is thus more reliable.

As a second approach, we also trained new random forest models using the training datasets of Park & Marcotte. The models that are summarized as $\text{ProteinPrompt}_{retrain}$ were trained similar to our original random forest model, i.e., we used the given protein pair orientation A-B from the training data file, but also the reversed orientation B-A and two combinations where one of both proteins has the inverted amino acid sequence: A-$B_{inv}$ and B-$A_{inv}$. This enhancement of the training space has small but consistent positive effects on the models.

Table S7 shows the results for the 'human balanced' dataset, S8 for 'human random', S9 for 'yeast balanced', S10 for 'yeast random'. The first thing to notice is that the accuracy for the human test sets decreases significantly when the random forest model is trained on the Park & Marcotte datasets. However, the achieved AUROC (area under the roc curve) and AUPRC (area under the precision recall curve) values are still highly comparable to the more powerful competitors.

This clearly suggests that our presented combination of autocorrelated biochemical residue scales and a random forest model is as good as other methodological approaches used to predict protein-protein interactions. The true benefit that clearly elevates ProteinPrompt comes with the extensive training and testing datasets that we collected and curated. Here, we could show that collecting as much training data as possible in combination with a rigorous cleaning can boost the prediction accuracy.

The decrease in accuracy for yeast in the normal version of ProteinPrompt, reveals a limitation in generalizability to distant organisms. This is not too surprising since learning methods are generally valid only for the training data domain. We could show that the accuracies increase when models are trained with yeast-specific data.

| | |
|---|---|
| C1 | Both proteins in the test pair are found in the training set |
| C2 | Only one protein in the test pair is found in the training set |
| C3 | Neither protein in the test pair is found in the training set |

Table S6: Definitions of C1, C2, C3 from [1]

| | AUROC | | | | AUPRC | | | |
|---|---|---|---|---|---|---|---|---|
| Method | CV | C1 | C2 | C3 | CV | C1 | C2 | C3 |
| ProteinPrompt$_{retrain}$ | 0.63 ± 0.01 | 0.64 ± 0.01 | 0.59 ± 0.01 | 0.54 ± 0.01 | 0.67 ± 0.01 | 0.69 ± 0.01 | 0.51 ± 0.03 | 0.54 ± 0.02 |
| ProteinPrompt | 0.87 ± 0.01 | 0.88 ± 0.01 | 0.77 ± 0.01 | 0.82 ± 0.01 | 0.90 ± 0.00 | 0.90 ± 0.01 | 0.83 ± 0.01 | 0.86 ± 0.01 |
| MMI+NMBAC | 0.61 ± 0.01 | 0.62 ± 0.01 | 0.57 ± 0.02 | 0.53 ± 0.01 | 0.64 ± 0.01 | 0.65 ± 0.01 | 0.58 ± 0.02 | 0.53 ± 0.01 |
| MMI | 0.61 ± 0.01 | 0.62 ± 0.01 | 0.57 ± 0.01 | 0.53 ± 0.01 | 0.64 ± 0.01 | 0.65 ± 0.01 | 0.58 ± 0.01 | 0.53 ± 0.01 |
| NMBAC | 0.59 ± 0.01 | 0.60 ± 0.01 | 0.56 ± 0.01 | 0.52 ± 0.02 | 0.62 ± 0.01 | 0.63 ± 0.01 | 0.56 ± 0.01 | 0.52 ± 0.01 |
| M1 | 0.64 ± 0.01 | 0.65 ± 0.01 | 0.61 ± 0.01 | 0.57 ± 0.02 | 0.66 ± 0.01 | 0.67 ± 0.01 | 0.61 ± 0.02 | 0.56 ± 0.02 |
| M2 | 0.59 ± 0.01 | 0.60 ± 0.01 | 0.06 ± 0.01 | 0.57 ± 0.02 | 0.60 ± 0.01 | 0.61 ± 0.01 | 0.60 ± 0.01 | 0.55 ± 0.01 |
| M3 | 0.55 ± 0.01 | 0.56 ± 0.01 | 0.54 ± 0.01 | 0.51 ± 0.01 | 0.60 ± 0.01 | 0.61 ± 0.01 | 0.55 ± 0.01 | 0.51 ± 0.01 |
| M4 | 0.56 ± 0.01 | 0.56 ± 0.01 | 0.54 ± 0.01 | 0.52 ± 0.01 | 0.54 ± 0.01 | 0.54 ± 0.01 | 0.53 ± 0.01 | 0.52 ± 0.01 |
| M5 | 0.59 ± 0.01 | 0.60 ± 0.01 | 0.56 ± 0.01 | 0.53 ± 0.01 | 0.63 ± 0.01 | 0.64 ± 0.01 | 0.57 ± 0.01 | 0.53 ± 0.01 |
| M7 | 0.55 ± 0.01 | 0.55 ± 0.02 | 0.53 ± 0.01 | 0.53 ± 0.02 | 0.55 ± 0.01 | 0.55 ± 0.01 | 0.53 ± 0.01 | 0.54 ± 0.02 |

Table S7: Comparison of the methods defined in [1, 2] with ProteinPrompt using the test datasets 'human balanced' provided by the Marcotte lab [3].

| | AUROC | | | | AUPRC | | | |
|---|---|---|---|---|---|---|---|---|
| Method | CV | C1 | C2 | C3 | CV | C1 | C2 | C3 |
| ProteinPrompt$_{retrain}$ | 0.83 ± 0.01 | 0.83 ± 0.01 | 0.59 ± 0.01 | 0.55 ± 0.02 | 0.84 ± 0.01 | 0.84 ± 0.01 | 0.60 ± 0.01 | 0.55 ± 0.02 |
| ProteinPrompt | 0.87 ± 0.01 | 0.87 ± 0.01 | 0.77 ± 0.01 | 0.82 ± 0.01 | 0.90 ± 0.00 | 0.90 ± 0.01 | 0.83 ± 0.01 | 0.86 ± 0.01 |
| MMI+NMBAC | 0.82 ± 0.01 | 0.82 ± 0.01 | 0.60 ± 0.01 | 0.57 ± 0.02 | 0.83 ± 0.01 | 0.83 ± 0.01 | 0.60 ± 0.01 | 0.56 ± 0.02 |
| MMI | 0.81 ± 0.01 | 0.81 ± 0.01 | 0.59 ± 0.01 | 0.56 ± 0.02 | 0.82 ± 0.01 | 0.83 ± 0.01 | 0.59 ± 0.01 | 0.55 ± 0.01 |
| NMBAC | 0.81 ± 0.01 | 0.82 ± 0.01 | 0.60 ± 0.01 | 0.57 ± 0.02 | 0.83 ± 0.01 | 0.83 ± 0.01 | 0.60 ± 0.01 | 0.56 ± 0.02 |
| M1 | 0.81 ± 0.01 | 0.81 ± 0.01 | 0.61 ± 0.01 | 0.58 ± 0.03 | 0.82 ± 0.01 | 0.82 ± 0.01 | 0.60 ± 0.01 | 0.57 ± 0.03 |
| M2 | 0.85 ± 0.01 | 0.85 ± 0.01 | 0.60 ± 0.01 | 0.58 ± 0.02 | 0.85 ± 0.01 | 0.85 ± 0.01 | 0.60 ± 0.01 | 0.56 ± 0.02 |
| M3 | 0.67 ± 0.01 | 0.68 ± 0.01 | 0.56 ± 0.01 | 0.51 ± 0.01 | 0.68 ± 0.01 | 0.68 ± 0.01 | 0.57 ± 0.01 | 0.51 ± 0.01 |
| M4 | 0.77 ± 0.01 | 0.77 ± 0.01 | 0.57 ± 0.02 | 0.53 ± 0.02 | 0.77 ± 0.01 | 0.77 ± 0.01 | 0.56 ± 0.01 | 0.53 ± 0.02 |
| M5 | 0.81 ± 0.01 | 0.81 ± 0.01 | 0.59 ± 0.01 | 0.54 ± 0.02 | 0.82 ± 0.01 | 0.82 ± 0.01 | 0.59 ± 0.01 | 0.54 ± 0.02 |
| M6 | 0.76 ± 0.01 | 0.77 ± 0.01 | 0.64 ± 0.01 | 0.59 ± 0.02 | 0.79 ± 0.01 | 0.80 ± 0.01 | 0.67 ± 0.01 | 0.60 ± 0.02 |
| M7 | 0.56 ± 0.01 | 0.56 ± 0.01 | 0.53 ± 0.01 | 0.54 ± 0.03 | 0.56 ± 0.01 | 0.56 ± 0.01 | 0.53 ± 0.01 | 0.54 ± 0.02 |

Table S8: Comparison of the methods defined in [1, 2] with ProteinPrompt using the test datasets 'human random' provided by the Marcotte lab [3].

| | AUROC | | | | AUPRC | | | |
|---|---|---|---|---|---|---|---|---|
| Method | CV | C1 | C2 | C3 | CV | C1 | C2 | C3 |
| ProteinPrompt$_{retrain}$ | 0.67 ± 0.02 | 0.69 ± 0.01 | 0.59 ± 0.02 | 0.53 ± 0.02 | 0.69 ± 0.02 | 0.71 ± 0.02 | 0.59 ± 0.02 | 0.53 ± 0.02 |
| ProteinPrompt | 0.56 ± 0.01 | 0.56 ± 0.01 | 0.53 ± 0.02 | 0.54 ± 0.02 | 0.55 ± 0.01 | 0.55 ± 0.01 | 0.53 ± 0.02 | 0.54 ± 0.03 |
| MMI+NMBAC | 0.65 ± 0.02 | 0.66 ± 0.02 | 0.60 ± 0.02 | 0.55 ± 0.02 | 0.67 ± 0.02 | 0.68 ± 0.02 | 0.60 ± 0.02 | 0.55 ± 0.02 |
| MMI | 0.64 ± 0.02 | 0.65 ± 0.01 | 0.60 ± 0.02 | 0.55 ± 0.02 | 0.66 ± 0.02 | 0.68 ± 0.01 | 0.60 ± 0.02 | 0.54 ± 0.02 |
| NMBAC | 0.63 ± 0.02 | 0.64 ± 0.02 | 0.59 ± 0.02 | 0.54 ± 0.03 | 0.65 ± 0.02 | 0.66 ± 0.02 | 0.59 ± 0.02 | 0.54 ± 0.02 |
| M1 | 0.64 ± 0.01 | 0.64 ± 0.01 | 0.62 ± 0.02 | 0.57 ± 0.04 | 0.65 ± 0.01 | 0.66 ± 0.01 | 0.62 ± 0.02 | 0.57 ± 0.03 |
| M2 | 0.61 ± 0.01 | 0.61 ± 0.02 | 0.62 ± 0.02 | 0.58 ± 0.03 | 0.62 ± 0.01 | 0.61 ± 0.02 | 0.62 ± 0.02 | 0.57 ± 0.03 |
| M3 | 0.55 ± 0.02 | 0.56 ± 0.02 | 0.55 ± 0.02 | 0.51 ± 0.01 | 0.59 ± 0.01 | 0.60 ± 0.01 | 0.56 ± 0.02 | 0.51 ± 0.02 |
| M4 | 0.55 ± 0.02 | 0.55 ± 0.02 | 0.54 ± 0.02 | 0.52 ± 0.01 | 0.53 ± 0.02 | 0.53 ± 0.01 | 0.54 ± 0.02 | 0.52 ± 0.02 |
| M5 | 0.60 ± 0.02 | 0.60 ± 0.01 | 0.55 ± 0.02 | 0.52 ± 0.02 | 0.61 ± 0.02 | 0.62 ± 0.01 | 0.55 ± 0.02 | 0.52 ± 0.02 |
| M7 | 0.55 ± 0.02 | 0.54 ± 0.01 | 0.54 ± 0.02 | 0.53 ± 0.02 | 0.55 ± 0.02 | 0.55 ± 0.01 | 0.54 ± 0.02 | 0.53 ± 0.02 |

Table S9: Comparison of the methods defined in [1, 2] with ProteinPrompt using the test datasets 'yeast balanced' provided by the Marcotte lab [3].

| | AUROC | | | | AUPRC | | | |
|---|---|---|---|---|---|---|---|---|
| Method | CV | C1 | C2 | C3 | CV | C1 | C2 | C3 |
| ProteinPrompt$_{retrain}$ | 0.82 ± 0.01 | 0.82 ± 0.01 | 0.58 ± 0.02 | 0.56 ± 0.03 | 0.84 ± 0.01 | 0.84 ± 0.01 | 0.59 ± 0.02 | 0.56 ± 0.02 |
| ProteinPrompt | 0.56 ± 0.01 | 0.56 ± 0.01 | 0.53 ± 0.02 | 0.54 ± 0.02 | 0.55 ± 0.01 | 0.55 ± 0.01 | 0.53 ± 0.02 | 0.54 ± 0.03 |
| MMI+NMBAC | 0.82 ± 0.02 | 0.82 ± 0.01 | 0.62 ± 0.02 | 0.61 ± 0.02 | 0.84 ± 0.01 | 0.84 ± 0.01 | 0.64 ± 0.02 | 0.62 ± 0.02 |
| MMI | 0.82 ± 0.01 | 0.82 ± 0.01 | 0.62 ± 0.02 | 0.60 ± 0.02 | 0.84 ± 0.02 | 0.84 ± 0.01 | 0.64 ± 0.02 | 0.61 ± 0.02 |
| NMBAC | 0.82 ± 0.01 | 0.82 ± 0.01 | 0.61 ± 0.02 | 0.60 ± 0.03 | 0.83 ± 0.01 | 0.83 ± 0.01 | 0.63 ± 0.03 | 0.60 ± 0.03 |
| M1 | 0.82 ± 0.01 | 0.82 ± 0.01 | 0.61 ± 0.02 | 0.58 ± 0.03 | 0.83 ± 0.02 | 0.83 ± 0.01 | 0.62 ± 0.02 | 0.58 ± 0.03 |
| M2 | 0.83 ± 0.01 | 0.84 ± 0.01 | 0.60 ± 0.02 | 0.59 ± 0.03 | 0.84 ± 0.02 | 0.85 ± 0.01 | 0.61 ± 0.02 | 0.58 ± 0.03 |
| M3 | 0.66 ± 0.02 | 0.67 ± 0.01 | 0.56 ± 0.02 | 0.51 ± 0.01 | 0.66 ± 0.02 | 0.66 ± 0.01 | 0.56 ± 0.02 | 0.51 ± 0.02 |
| M4 | 0.76 ± 0.02 | 0.76 ± 0.02 | 0.57 ± 0.02 | 0.54 ± 0.03 | 0.76 ± 0.02 | 0.76 ± 0.02 | 0.58 ± 0.02 | 0.54 ± 0.03 |
| M5 | 0.80 ± 0.02 | 0.80 ± 0.01 | 0.57 ± 0.01 | 0.55 ± 0.02 | 0.81 ± 0.02 | 0.81 ± 0.02 | 0.59 ± 0.02 | 0.55 ± 0.02 |
| M6 | 0.75 ± 0.02 | 0.75 ± 0.02 | 0.59 ± 0.04 | 0.55 ± 0.03 | 0.79 ± 0.02 | 0.79 ± 0.02 | 0.64 ± 0.03 | 0.55 ± 0.04 |
| M7 | 0.58 ± 0.02 | 0.58 ± 0.01 | 0.54 ± 0.02 | 0.53 ± 0.02 | 0.60 ± 0.02 | 0.60 ± 0.02 | 0.55 ± 0.02 | 0.53 ± 0.02 |

Table S10: Comparison of the methods defined in [1, 2] with ProteinPrompt using the test datasets 'yeast random' provided by the Marcotte lab [3].

[3] Training and test datasets for human and yeast: http://www.marcottelab.org/differentialGeneralization/
